# Enhanced Sampling of Protein Conformational Transitions via Dynamically Optimized Collective Variables

**DOI:** 10.1101/390278

**Authors:** Z. Faidon Brotzakis, Michele Parrinello

## Abstract

Protein conformational transitions often involve many slow degrees of freedom. Their knowledge would give distinctive advantages since it provides chemical and mechanistic insight and accelerates the convergence of enhanced sampling techniques that rely on collective variables. In this study, we implemented a recently developed variational approach to conformational dynamics metadynamics to the conformational transition of the moderate size protein, L99A T4 Lysozyme. In order to find the slow modes of the system we combined data coming from NMR experiments as well as short MD simulations. A Metadynamics simulation based on these information reveals the presence of two intermediate states, at an affordable computational cost.

## 1 Introduction

The L99A variant of the T4 Lysozyme (L99A T4L) protein exhibits two metastable states, G and E (see Figure 1a,b). The former is the high populated ground state while the latter is a low populated excited state. The interconversion between the two takes place on the timescale of miliseconds^1,2^. In the G state a large hydrophobic cavity (150 Å^3^) is formed, in contrast with state E in which the hydrophobic cavity is filled by the F114 residue^1–3^. Even more intriguingly L99A T4L is druggable and several crystal structures of the holo form have been reported^3,4^. It is therefore a prototypical example of protein plasticity.

**Figure 1:**
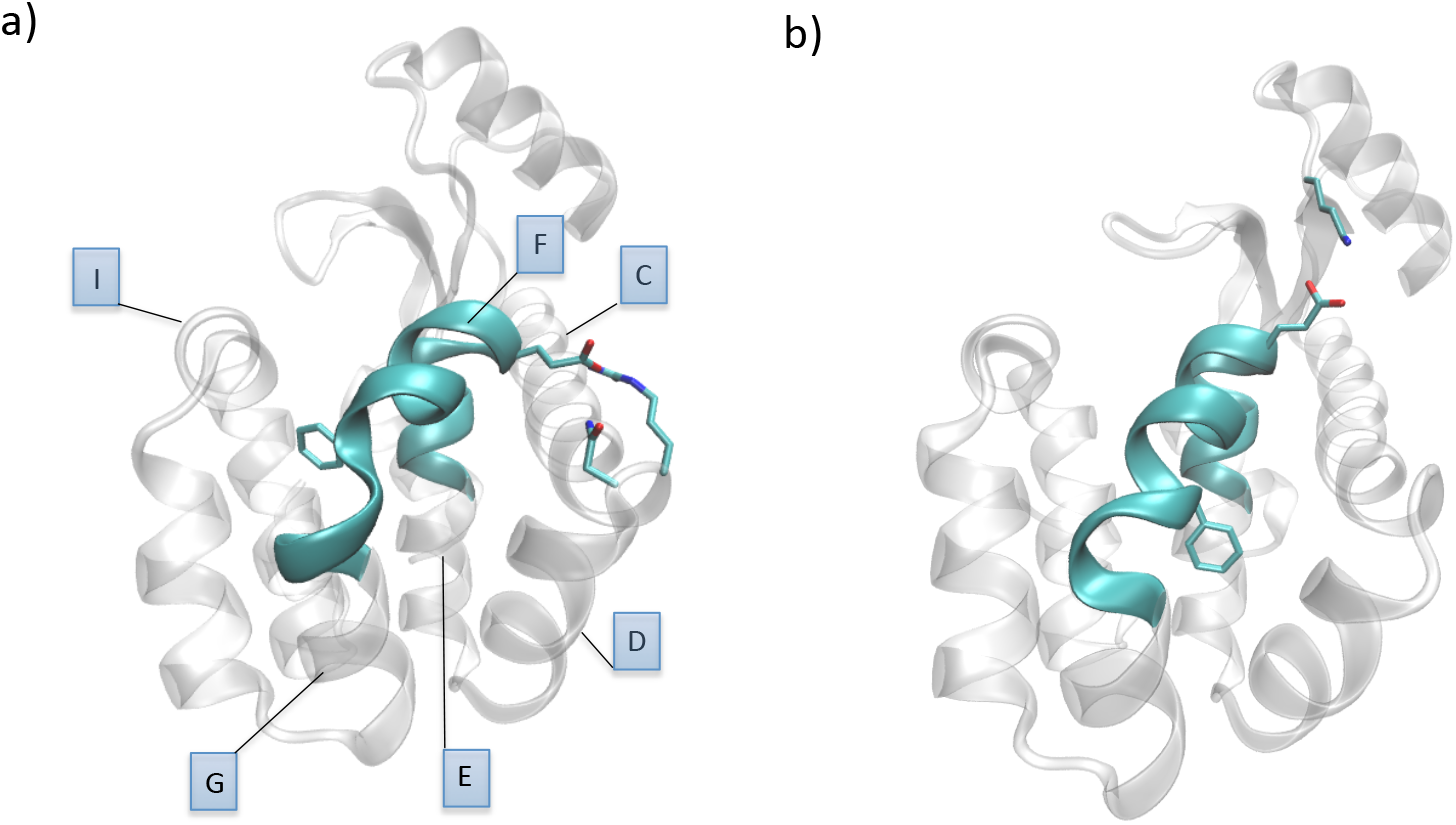
Representation of the L99A T4 Lysozyme in a) state G and, b) in state E. Note that in cyan is highlighted the region of aminoacids showing an increased chemical shift signal^1^. In licorice representation we show the important salt, bridge formed between residues E108 and R80,N81 in state G and between residues E108 and K35 in state E. Surrounding helices are labelled in blue.

Since the G to E transformation takes place on a milisecond timescale a direct Molecular Dynamics (MD) simulation is outside our present computational capability, especially if one wants to gather sufficient statistics. One way to tackle this timescale problem is to use enhanced sampling methods. One such method is Metadynamics (MetaD) that belongs to the class of enhanced sampling methods that use Collective Variables (CVs) to describe the slow modes involved in the transition of interest. In a remarkable application of MetaD, Wang et al made use of Path Collective Variables Metadynamics (Path CV-MetaD) and were able to discover an intermediate metastable state that is capable of accommodating a ligand.

In this paper we want to complement the effort of Wang et al by approaching the same problem from a different angle. Our purpose is to reveal what are the slow modes that are to be associated with the G to E transition. This will also assess the ability of recently developed methods of obtaining such modes in a relatively more complex scenario such as that of a real protein. Our approach will be based on Variational Approach to Conformational dynamics Metadynamics (VAC-MetaD). In VAC-MetaD one first performs a short MetaD run with non optimal CVs that later uses to derive more efficient CVs. Since CV efficiency is strongly related to how well they describe the physical process one gains not only efficiency but also insight. Using these CVs we not only rediscover the on pathway intermediate state of Ref.^5^, but also we find a new one.

## 2 Methods

### 2.1 Metadynamics

Metadynamics (MetaD) is a well known enhanced sampling method for studying rare events^6–8^. In MetaD one speeds up sampling by adding a history-dependent bias constructed by depositing Gaussian kernels on selected degrees of freedom, known as collective variables (CVs).

In well-tempered Metadynamics^9^ this goal is achieved by periodically adding a bias that is built according to the iterative procedure

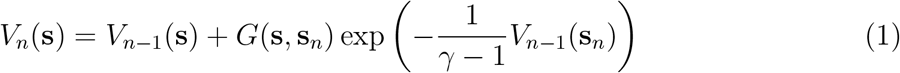

where V_*n*_(s) is the total bias deposited at iteration *n* and is obtained by adding to V_*n*–1_ a Gaussian term *G*(s, s_*n*_) centered on the present value of the CV and scaled by the factor 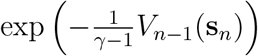. This factor makes the height of the added Gaussian decay with time. The bias factor *γ* determines the rate with which the added bias decreases and regulates the s fluctuations. In this way, the MetaD potential allows the system to explore the FES, escaping the free-energy minima and crossing large energy barriers at an affordable computational cost.

An advantage of well-tempered MetaD is that, after a transient period, the simulation enters a quasi-stationary limit in which the expectation value of any observable 〈*O*(**R**)〉^10^, can be estimated as running average according to:

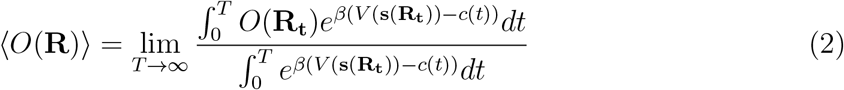

where **R**_*t*_ are atomic positions at time *t, β* is the reciprocal of the temperature multiplied by the Boltzmann constant k_*B*_ and c(t) is a time dependent offset defined as:

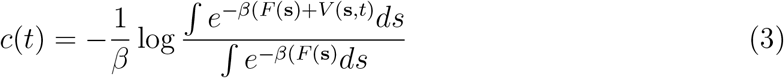

where F(s) is the free energy as a function of the CV. In this way, one can reweight to Boltzmann averages of any variables even if they are not included in the biased CV.

### 2.2 Variational Approach to Conformational dynamics Metadynamics

In MetaD the cost of reconstructing the free energy grows exponentially with the number of CVs, therefore, the number of CVs should be kept as small as possible. This condition prevent us from describing all relevant degrees of freedom of the system. This limitation is particularly restrictive in proteins where a slow process often involves crossing multiple small barriers, involving several degrees of freedom. In 2017 McCarty and Parrinello developed the Variational Approach to Conformational dynamics in Metadynamics (VAC-MetaD)^11^ that allows combining several degrees of freedom into a number of CVs that are linear expansions of several descriptors. So far this approach has shed light into chemical reactions and the folding of small peptides^12,13^.

Here we review shortly the necessary steps for applying this procedure. Initially one identifies a set of descriptors *d_k_*(**R**) that are important for the process under study. Then one searches for the best CV s_*i*_(**R**) that can be expressed as a linear combination of d_*k*_(**R**):

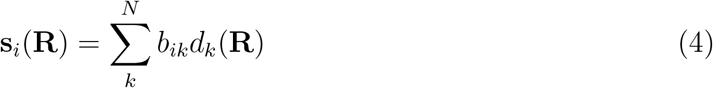

This is achieved by a variant of Time-lagged Independent Conformational Analysis (TICA) developed in Noe’s group^14,15^. In this framework, one computes the time-lagged covariance matrix C(*τ*) defined as:

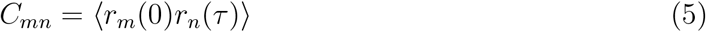

where r_*k*_(*τ*)=d_*k*_(*τ*)-〈*d_k_*〉. The expansion coefficients are determined by solving the eigenvalue equation:

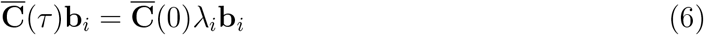

where 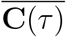 is the time lagged covariance matrix, 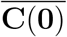 is the same matrix at time zero, *λ_i_* is the *i*th eigenvalues and **b**_*i*_ is the *i*th vector of coefficients of the CV s_*i*_. In Ref.^14,15^, one starts from the assumptuon that a trajectory that already crosses the barriers several times is available. McCarty and Parrinello^11^ showed that an analogous procedure can be applied to a MetaD trajectory, if the MetaD timescale was properly rescaled:

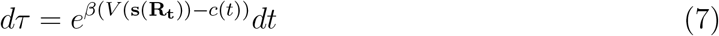

For the purpose of studying a protein conformational transitions, the eigenvectors of the eigenvalue equation obtained from a MetaD trajectory, correspond to the slowest decaying modes, and can be used as CVs in the production MetaD simulation^11,16^.

### 2.3 Selecting the set of descriptors

In the application of VAC it is essential to choose the right set of descriptors. In this choice we shall be guided mostly by experimental facts. NMR chemical shifts indicate that residues 101 to 118, whose position is highlighted in Figure 1 are mobile on the milisecond timescale. We choose to express their motion in terms of descriptors related to these residues such as dihedral angles and contacts, as detailed in Table S1 and Figure S1. The rest of the descriptors come from an analysis of two short MD simulations in state G and E. These simulations showed that a salt bridge away from the hydrophobic cavity between E108 with R80, N81 and E108 with K35 did change in the two states. For this reason we expressed these two possibilities in terms of two coordination number descriptors.

### 2.4 Protocol

We make use of the CHARMM27 and TIP3P forcefields for the protein and water respectively^17,18^. All our MetaD simulations initiate from state G of L999A T4L, obtained from X-ray crystal structure (PDB ID: 3DMV). For more information about the system preparation we refer the reader to the SI. Briefly, we performed a 350 ns NPT MetaD simulation at 310 K and atmospheric pressure, starting from state G and biasing a suboptimal CV constructed as an equal weights linear combination of the descriptors mentioned in the subsection IIB. This led to the Free Energy Surface (FES) depicted in Figure 3a where only the G and E states are explored. After that we analysed this trajectory as described in subsection IIB and obtained the eigenvalues shown in Figure 2a. It can be seen that the two top most eigenvalues decay more slowly than the others. We thus chose two corresponding eigenvectors as CVs, henceforth abbreviated s_1_ and s_2_. An analysis of these eigenvectors shows that not all 21 descriptors are relevant. As expected, the descriptors 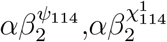 related to the swing of phenylalanine F114 from states G to E are relevant. Then, we find that the H-bond contacts descriptors (CN_1761–1803_,CN_1789–1751_,CN_1817–1768_,CN_1788–1828_) related to the rewinding of the F helix in going from G to E, play a role. Also relevant is the descriptor 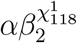 depicting the side-chain rotation of the hydrophobic residue L_118_. Finally, the relevance of the salt bridges between residues E108 and R80, N81 and residues E108 and K35 (CN_*SBG*_ and CN_*SBE*_) is apparent. It is also to be noted a tendency of the CN_*SBG*_ and CN_*SBE*_ to be in anti-phase. For a detailed explanation of the descriptors the reader can go to Table S1 and Figure S1.

**Figure 2:**
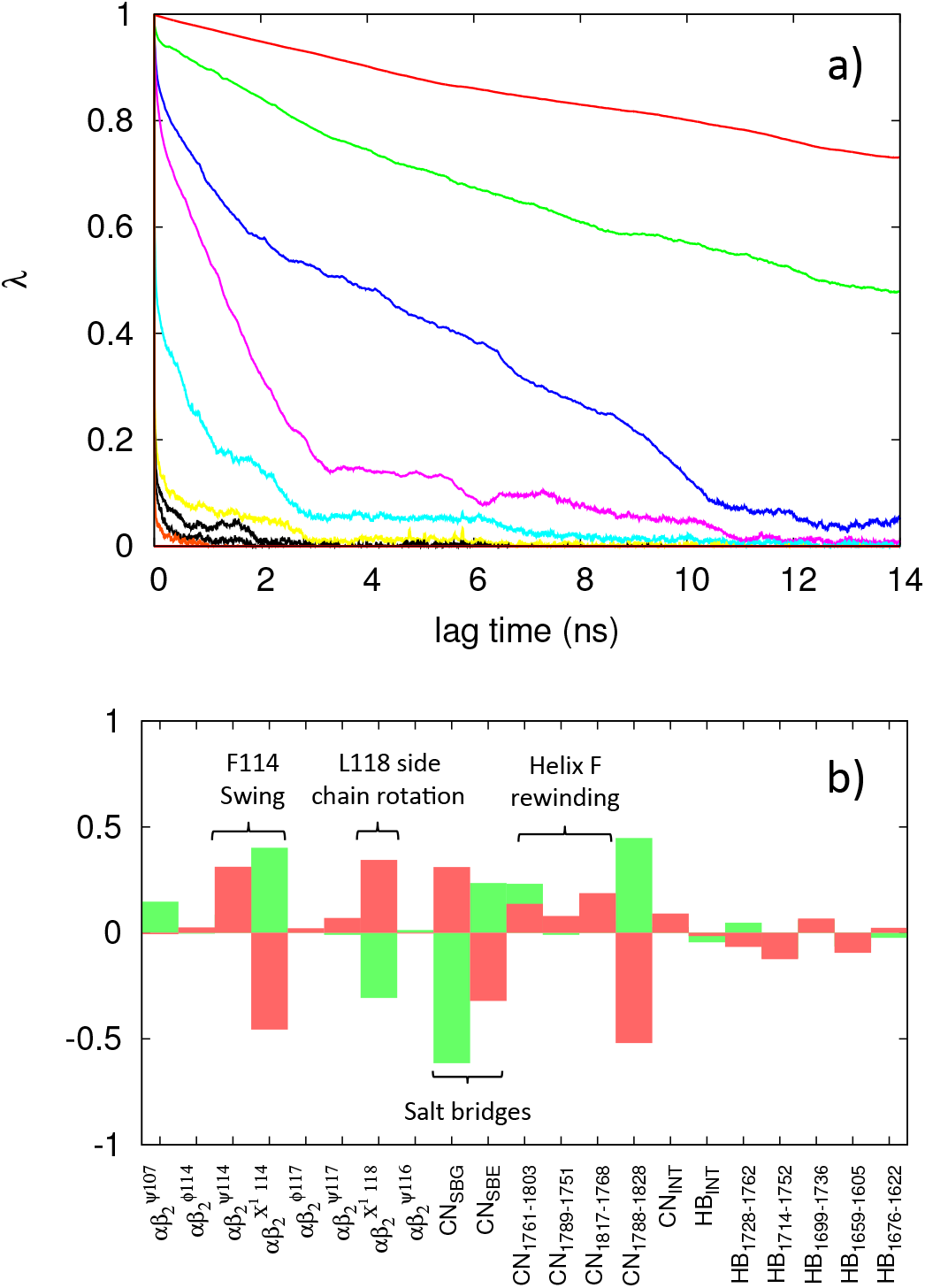
a) Eigenvalues decay as a function of lag time and b) coefficients of the set of descriptors for the two eigenvectors associated with the two first slowly decaying eigenvalues (red and green respectively).

**Figure 3:**
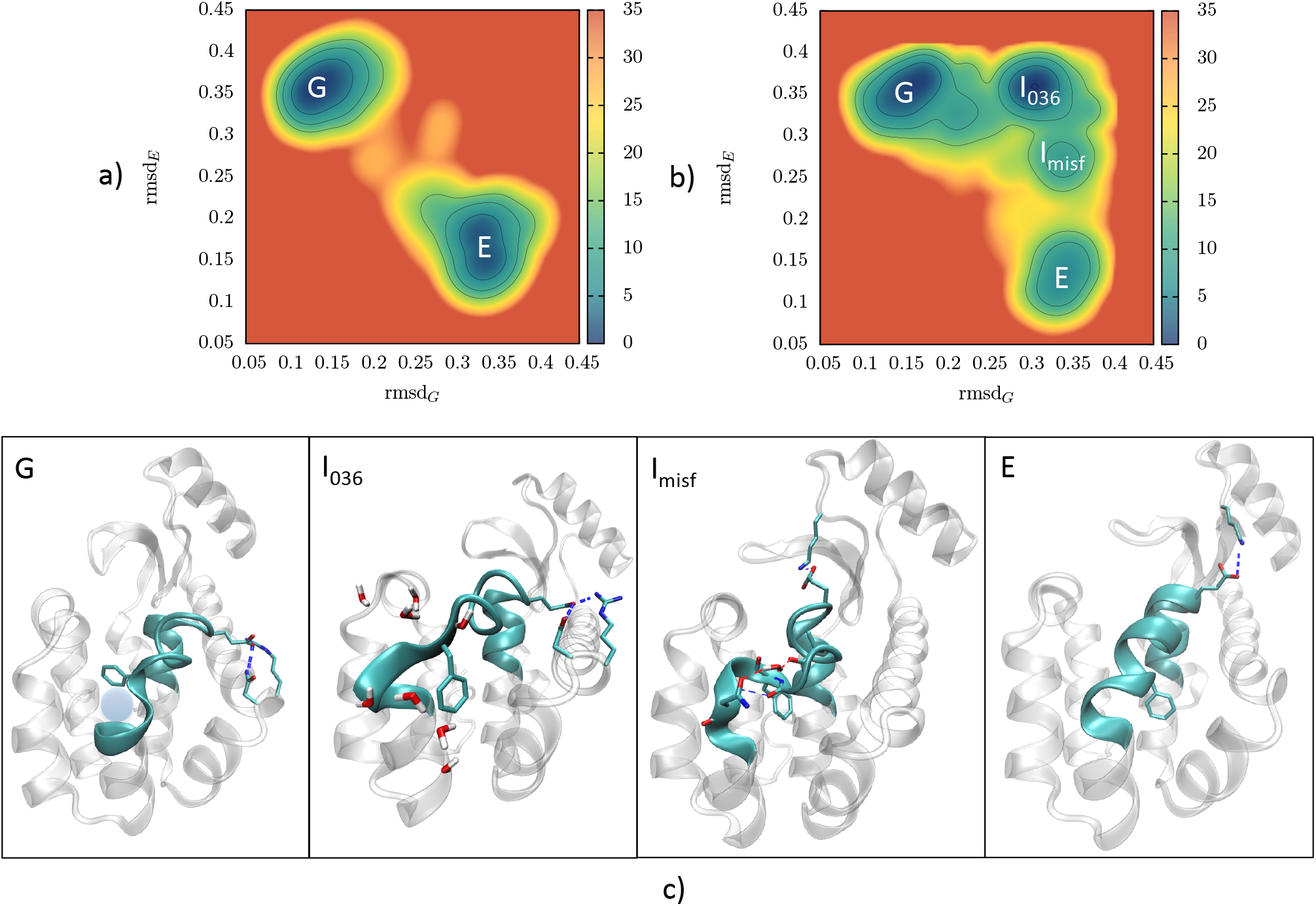
FES in kJ/mol as a function of the RMSD_*G*_ and RMSD_*E*_ for a) the MetaD and b) the VAC-MetaD simulation, c) States G, I_036_, I_*misf*_ and E explored during the VAC-MetaD simulation. In shaded purple is depicted schematically the position of the hydrophobic cavity of state G.

## 3 Results

Using the VAC-MetaD derived CVs s_1_ and s_2_ we performed a new 800 ns MetaD simulation, henceforth abbreviated as VAC-MetaD. Not only does the system go reversibly directly from G to E but remarkably it also follows a different route by passing through two other metastable intermediates, shown in Figure 3b. One is the previously found I_036_, the other is a newly found metastable intermediate, abbreviated I_*misf*_. Both metastable intermediates were verified to be stable after running unbiased simulations of 20 ns for each of the states.

### 3.1 Conformational free energies

In order to give quantitative insight on the population of the for states, we compute the free energy differences between the states in both MetaD and VAC-MetaD simulations. Regarding the technical aspects of this procedure we refer the interested reader to the SI.

The MetaD performed with suboptimal CVs gives a free energy difference between states G and E of 0.2 ± 2kJ/mol. This value is not close to the experimentally known value of 7.35 ± 0.8 kJ/mol^2^ ns which we attribute to the fact that the CV is suboptimal and the simulation is not fully converged. However, the free energy difference between G and E obtained from the VAC-MetaD simulations is 5.2 ± 2.0 kJ/mol, which is in excellent agreement with the experiment, thus proving the quality of these CVs. The free energy difference between G and I_036_ is found to be 5.6 ± 4.4 kJ/mol and the one between G and I_*misf*_ is 14.8 ± 4.6 kJ/mol. Notably, state I_*misf*_ is found to be much higher in energy compated to state G. We attribute this to the fact that in I_*misf*_ the protein assumes a misfolded conformation.

### 3.2 Description of the states

Here, we give a brief description of the four states found and we refer to the interested reader to the Supporting Information.

#### State G

Our VAC-MetaD simulation captures well the minimum of state G as shown in Figure 3b and Figure 3c,G. In agreement with the crystal structure of state G, we verify that F114 residue is turned with an angle *ψ* at 49°, and buried inside the cavity, while making hydrophobic contacts with aminoacids of helices G, E, I^1,3^. Another characteristic feature of this state is the type of winding of the first part of helix G (residues 116-121) with a kink at the position of residues 112-117. Note also the presence of the salt bridge between aminoacids E108 with R80 and N81 that helps stabilization of this state although distant from hydrophobic cavity. The hydrophobic cavity highlighted in Figure 3c,G. is made inaccessible by the presence of phenylalanine F114.

#### State I_036_

Some of the most important features of this state are the rotation of F114 that is now exposed to the solvent, the partial unfolding of the last part of helix F (residues 113-115), and the permanence of the salt bridge between E108 with R80 and N81. The new F114 position is now stabilized by hydrophobic contacts with G and D helices. The formation of a cavity is able to accommodate a ligand, its shape and size can be seen in Figure 4a.

**Figure 4:**
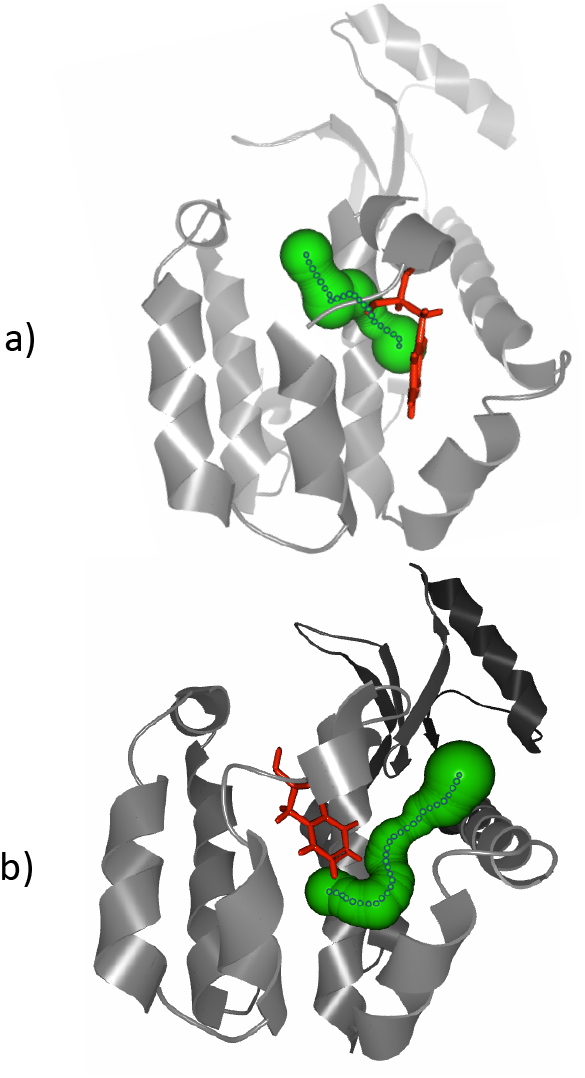
The most probable tunnel formed in states a) I_036_, formed between helices F and I and b) I_*misf*_, formed between helices F,E and D.

#### State E

Before describing I_*misf*_ it is more illuminating to first describe the E state. Since I_*misf*_ can be looked at as a modification of the E state in the same way as I_036_ is best understood moving reference to G. In the E state the kink between helices G and F moves to the position of residues 115 to 120, and at the end of helix F hydrogen bonds between residues 111 and 116 are formed. Also, here there is a stabilizing salt bridge but now between E108 and K35.

#### State I_*misf*_

In this state a some H-bonds that stabilized helices G and F in state E are broken and now are replaced by new ones, a typical characteristic of misfolding. More precisely, direct H-bonds between F114 and S117 and a water mediated hydrogen bond between G113 and N116 are formed. Now F114 assumes an orientation intermediate between those that this residue has in G and E. The E108-K35 salt bridge present in state E is maintained. An elongated cavity that sits between the F, E and D helices is formed (see Figure 4b). Contrary to state I_036_ the diameter of this cavity is too small to allow a ligand insertion. The F114 is now in an intermediate position in the hydrophobic cavity between states G and E. Like in state E, there is a stabilizing salt bridge between E108 and K35.

## 4 Discussion

In this study we implemented for the first time the VAC-MetaD framework to a conformational transition of the moderate size protein, L99A T4 Lysozyme. Our aim was to obtain insight in the slow degrees of freedom, by looking at the slowly decaying eigenvectors of the VAC-MetaD analysis, and accelerate convergence using these dynamically optimized CVs and unravel the complex free energy landscape of this conformational transition.

Combining experimental knowledge from NMR and short MD simulations in states G and E and applying the VAC-MetaD framework to an initial MetaD trajectory we unravelled the slow degrees of freedom. In particular these involve, the swing motion of phenylalanine F114 in going from G to E, the H-bond contacts related to the rewinding of helix F in going from G to E, a side chain rotation of Leucine L118 in the hydrophobic cavity and finally, a salt bridge formation away from the hydrophobic cavity between residues E108 and K35 or residues E108 and R80 and N81. Convergence was much improved using the VAC-MetaD optimized CVs, compared to suboptimal CVs. The free energy difference between state G and E was found in very good agreement with experiments.

Strikingly, while the MetaD simulation only contained direct transitions between states G and E, the VAC-MetaD optimized CVs contained more information, and when biased where able to drive the system transitioning through an unprecedented route, passing through two other metastable intermediate states I_036_ and I_*misf*_. The first is a previously found metastable intermediate state able to accommodate a ligand, and the second a newly found state with a misfolded F helix, higher in energy and unable to accommodate a ligand.

This work, proves the power of VAC-MetaD in accelerating convergence of the free energy, exploring the underlying free energy landscape and providing chemical insight through the VAC-MetaD eigenvectors. We see VAC-MetaD analysis as a tool that achieves all three goals and is up to the user whether he pursuits all of them. For example, a VAC-MetaD analysis of an initial MetaD or MD trajectory, is already insightful giving acess to the slow degrees of freedom.

A downside of the VAC-MetaD is that of the initial trajectory necessary to to optimize the descriptor set CVs. Hence, we anticipate that faster methods such as Variational Enhanced Sampling and/or multiple walkers would help obtaining the initial trajectory faster.

## 5 Associated content

Supporting Information Available: Simulation details for MD and Metadynamics, free energy difference calculation details, free energy landscape identification, and table of descriptor details.

## 6 Acknowledgements

This research was supported by the VARMET European Union Grant ERC—2014—ADG—670227. Computational resources were provided by the Swiss National Supercomputing Centre (CSCS).

---

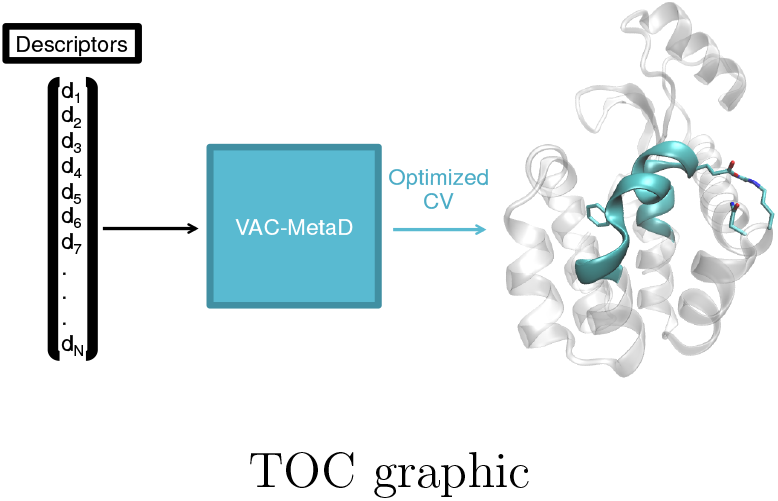
TOC graphic

## References

(1) Bouvignies, G.; Vallurupalli, P.; Hansen, D. F.; Correia, B. E.; Lange, O.; Bah, A.; Vernon, R. M.; Dahlquist, F. W.; Baker, D.; Kay, L. E. Solution structure of a minor and transiently formed state of a T4 lysozyme mutant. Nature 2011, 477, 111–117.

(2) Mulder, F. A. A.; Mittermaier, A.; Hon, B.; Dahlquist, F. W.; Kay, L. E. Studying excited states of proteins by NMR spectroscopy. Nat. Struct. Biol. 2001, 8, 932–935.

(3) Eriksson, A. E.; Baase, W. A.; Wozniak, J. A.; Matthews, B. W. A cavity-containing mutant of T4 lysozyme is stabilized by buried benzene. Nature 1992, 355, 371–373.

(4) Merski, M.; Fischer, M.; Balius, T. E.; Eidam, O.; Shoichet, B. K. Homologous ligands accommodated by discrete conformations of a buried cavity. Proc. Natl. Acad. Sci. 2015, 112, 5039–5044.

(5) Wang, Y.; Papaleo, E.; Lindorff-Larsen, K. Mapping transiently formed and sparsely populated conformations on a complex energy landscape. Elife 2016, 5, 1–35.

(6) Laio, A.; Parrinello, M. Escaping free-energy minima. Proc. Natl. Acad. Sci. U. S. A. 2002, 99, 12562–6.

(7) Barducci, A.; Bussi, G.; Parrinello, M. Well-Tempered Metadynamics: A Smoothly Converging and Tunable Free-Energy Method. Phys. Rev. Lett. 2008, 100, 1–4.

(8) Barducci, A.; Bonomi, M.; Parrinello, M. Metadynamics. Wiley Interdiscip. Rev. Comput. Mol. Sci. 2011, 1, 826–843.

(9) Bonomi, M.; Parrinello, M. Enhanced sampling in the well-tempered ensemble. Phys. Rev. Lett. 2010, 104, 1–4.

(10) Tiwary, P.; Dama, J. F.; Parrinello, M. A perturbative solution to metadynamics ordinary differential equation. J. Chem. Phys. 2015, 143, 1–4.

(11) Mccarty, J.; Parrinello, M.; Mccarty, J.; Parrinello, M. A variational conformational dynamics approach to the selection of collective variables in metadynamics A variational conformational dynamics approach to the selection of collective variables in metadynamics. J. Chem. Phys. 2017, 147, 204109–1.

(12) Piccini, G.; Polino, D.; Parrinello, M. Identifying Slow Molecular Motions in Complex Chemical Reactions. J. Phys. Chem. Lett. 2017, 8, 4197–4200.

(13) Yang, Y. I.; Parrinello, M. Refining Collective Coordinates and Improving Free Energy Representation in Variational Abstract Keywords I Introduction. J. Chem. Theory Comput. 2018, 14, 2889–2894.

(14) Pérez-hernández, G.; Paul, F.; Giorgino, T.; Fabritiis, G. D.; Noé, F. Identification of slow molecular order parameters for Markov model construction Identification of slow molecular order parameters for Markov. 2013, 015102–13.

(15) Noé, F.; Nüske, F. Copyright © by SIAM. Unauthorized reproduction of this article is prohibited. Copyright © by SIAM. Unauthorized reproduction of this article is prohibited. Multiscale Model. Simul. 2013, 11, 635–655.

(16) Sultan, M. M.; Pande, V. S. TICA-Metadynamics: Accelerating Metadynamics by Using Kinetically Selected Collective Variables. J. Chem. Theory Comput. Theory Comput. 2017, 13, 2440–2447.

(17) Jorgensen, W. L.; Madura, J. D. Solvation and Conformation of Methanol in Water. J. Am. Chem. Soc. 1983, 105, 1407–1413.

(18) Blanchet, C.; Pasi, M.; Zakrzewska, K.; Lavery, R.; Moakher, M.; Maddocks, J. H.; Petkeviciute, D.; Zakrzewska, K.; Nikolov, D. B.; Chen, H.; Halay, E. D.; Hoffmann, A.; Roeder, R. G.; Burley, S. K.; Chua, E. Y. D.; Vasudevan, D.; Davey, G. E.; Wu, B.; Davey, C. A.; Foloppe, N.; Mackerell, A. D.; Feig, M.; Brooks, C. L.; Bashford, D.; Bellott, M.; Dunbrack, R. L.; Evanseck, J. D.; Field, M. J.; Fischer, S.; Gao, J.; Guo, H.; Ha, S.; Joseph-McCarthy, D.; Kuchnir, L.; Kuczera, K.; Lau, F. T. K.; Mattos, C.; Michnick, S.; Ngo, T.; Nguyen, D. T.; Prodhom, B.; Reiher, W. E.; Roux, B.; Schlenkrich, M.; Smith, J. C.; Stote, R.; Straub, J.; Watanabe, M.; Wiorkiewicz-Kuczera, J.; Yin, D.; Karplus, M.; Banavali, N. K.; Bjelkmar, P.; Larsson, P.; Cuendet, M. A.; Hess, B.; Lindahl, E. Implementation of the CHARMM force field in GRO-MACS: Analysis of protein stability effects from correction maps, virtual interaction sites, and water models. J. Chem. Theory Comput. 2010, 6, 459–466.

